# The Early Pregnancy Human Placenta Shares Hypomethylation Patterns Characteristic of Solid Tumors

**DOI:** 10.1101/098616

**Authors:** Akpéli V. Nordor, Djamel Nehar-Belaid, Sophie Richon, David Klatzmann, Dominique Bellet, Virginie Dangles-Marie, Thierry Fournier, Martin J. Aryee

## Abstract

**Background:** The placenta relies on phenotypes that are characteristic of cancer to successfully implant the embryo in the uterus during early pregnancy. Notably, it has to invade its host tissues, promote angiogenesis, while surviving hypoxia, and escape the immune system. Similarities in DNA methylation patterns between the placenta and cancers suggest that common epigenetic mechanisms may be involved in regulating these behaviors.

**Results:** We show here that megabase-scale patterns of hypomethylation distinguish first from third trimester chorionic villi in the placenta, and that these patterns mirror those that distinguish many tumors from corresponding normal tissues. We confirmed these findings in villous cytotrophoblasts isolated from the placenta and identified a time window at the end of the first trimester, when these cells come into contact with maternal blood as the likely time period for the methylome alterations. Furthermore, the large genomic regions affected by these patterns of hypomethylation encompass genes involved in pathways related to epithelial-mesenchymal transition (EMT), immune response and inflammation. Analyses of expression profiles corresponding to genes in these hypomethylated regions in colon adenocarcinoma tumors point to networks of differentially expressed genes previously implicated in carcinogenesis and placentogenesis, where nuclear factor kappa B (NF-kB) is a key hub.

**Conclusion:** Taken together, our results suggest the existence of epigenetic switches involving large-scale changes of methylation in the placenta during pregnancy and in tumors during neoplastic transformation. The characterization of such epigenetic switches might lead to the identification of biomarkers and drug targets in oncology as well as in obstetrics and gynecology.

## BACKGROUND

At the beginning of pregnancy, human placental trophoblasts rely on several phenotypes that are also hallmarks of cancer in order to implant the blastocyst containing the embryo in the uterus. After undergoing epithelial-mesenchymal transition, placental cells invade and migrate within the endometrium. They also promote angiogenesis, while surviving hypoxia, to establish fetal/maternal exchange of nutrients, gases and wastes. Moreover, as 50% of the placental genome has paternal origin, placental cells have to evade the maternal immune system [1–4].

Studies on DNA methylation in cancer cells and placental cells have highlighted similarities in their epigenetic landscapes which are characterized by a widespread hypomethylation throughout the genome and focal hypermethylation at CpG islands, including at promoters of tumor suppressor genes [5, 6]. A recent study also linked the observation of genes specifically expressed in both the placenta and various tumors to hypomethylation [7]. However, to our knowledge, no study has yet directly compared the genome-wide DNA methylation alterations in placenta and cancers. We hypothesized that there may be parallels between the patterns of methylation that distinguish the early pregnancy placenta from the late pregnancy placenta and those that distinguish tumors from corresponding normal tissues, and that such similar epigenetic patterns may contribute to the regulation of shared cancer/placenta phenotypes.

Therefore, we conducted a genome-wide comparison of DNA methylation changes in placental tissues during pregnancy and in 13 types of tumor tissues during neoplastic transformation. We used publically available placenta and cancer DNA methylation data, complemented by methylome data generated from villous cytotrophoblasts isolated from placental tissues. All DNA methylation profiling was performed using the Illumina HumanMethylation450 (450k) microarray platform, thereby facilitating comparisons across datasets. We also investigated links between cancer/placenta patterns of methylation and gene expression profiles in cancer.

We report here that the early pregnancy human placenta is hypomethylated compared to that later in pregnancy. This hypomethylation is organized into megabase-scale domains that frequently overlap with hypomethylated block regions observed in solid tumors [8, 9]. These hypomethylated blocks encompass genes involved in mechanisms related to hallmark cancer phenotypes.

## RESULTS

### Placental methylome remodeling during pregnancy involves widespread methylation gains in CpG-poor genomic regions

To identify epigenomic features specific to the early pregnancy placenta, we first examined DNA methylation changes that occur between first and third trimester chorionic villi samples using publically available data (Gene Expression Omnibus (GEO), <u>GSE44667</u>) [10] (Fig. 1a; Table S1). The chorionic villus represents the structural and functional unit that projects from the fetal placenta to invade the maternal uterine lining. The villous core, made of fibroblasts, mesenchymal cells, Hofbauer cells and fetal-placental vessels, is covered by layers of villous cytotrophoblasts and syncytiotrophoblasts, which are in direct contact with maternal blood [11]. DNA methylation was measured on the 450k array and quantified using Beta values. We computed the difference in methylation between first trimester (6-10 weeks of gestation (WG), n = 5) and third trimester (32-39 WG, n = 10) chorionic villi samples. We found that while there were many alterations at CpGs distal from CpG islands with 36% of CpGs affected in open-sea regions (> 4 kb from the nearest CpG island [12]), there were relatively few methylation changes at CpG islands themselves (11%) (Fig. 1b; Table S2, Figure S1 and Figure S2). The majority (72%) of changes in open-sea regions represent hypomethylation (methylation difference < - 0.05, FDR q-value < 0.05) in first *vs*. third trimester chorionic villi. In contrast, methylation differences at CpG islands, in addition to being much less frequent, were equally balanced between hypermethylation (53%) and hypomethylation (47%) (Fig. 1c). We next compared placental methylomes to those of tumors profiled on the 450k array by The Cancer Genome Atlas (TCGA, http://cancergenome.nih.gov/) for 13 cancer types (4,115 primary tumor and 460 matched normal tissue samples) (Fig 1a; Table S1). As expected, cancer tissues displayed large changes at both CpG islands (32% altered) and at CpGs distal from islands (45% altered) as illustrated with colon adenocarcinoma (Fig. 1b; Table S2, Figure S1 and Figure S2). Hypomethylation represents 75% of changes in open sea regions, while hypermethylation represents 78% of changes at CpG islands (Fig. 1c). Taken together, these results suggest open-sea hypomethylation as a common epigenetic feature of early placental cells and neoplastic cells, and that this hypomethylation begins to resolve as pregnancy progresses.

**Fig 1.**
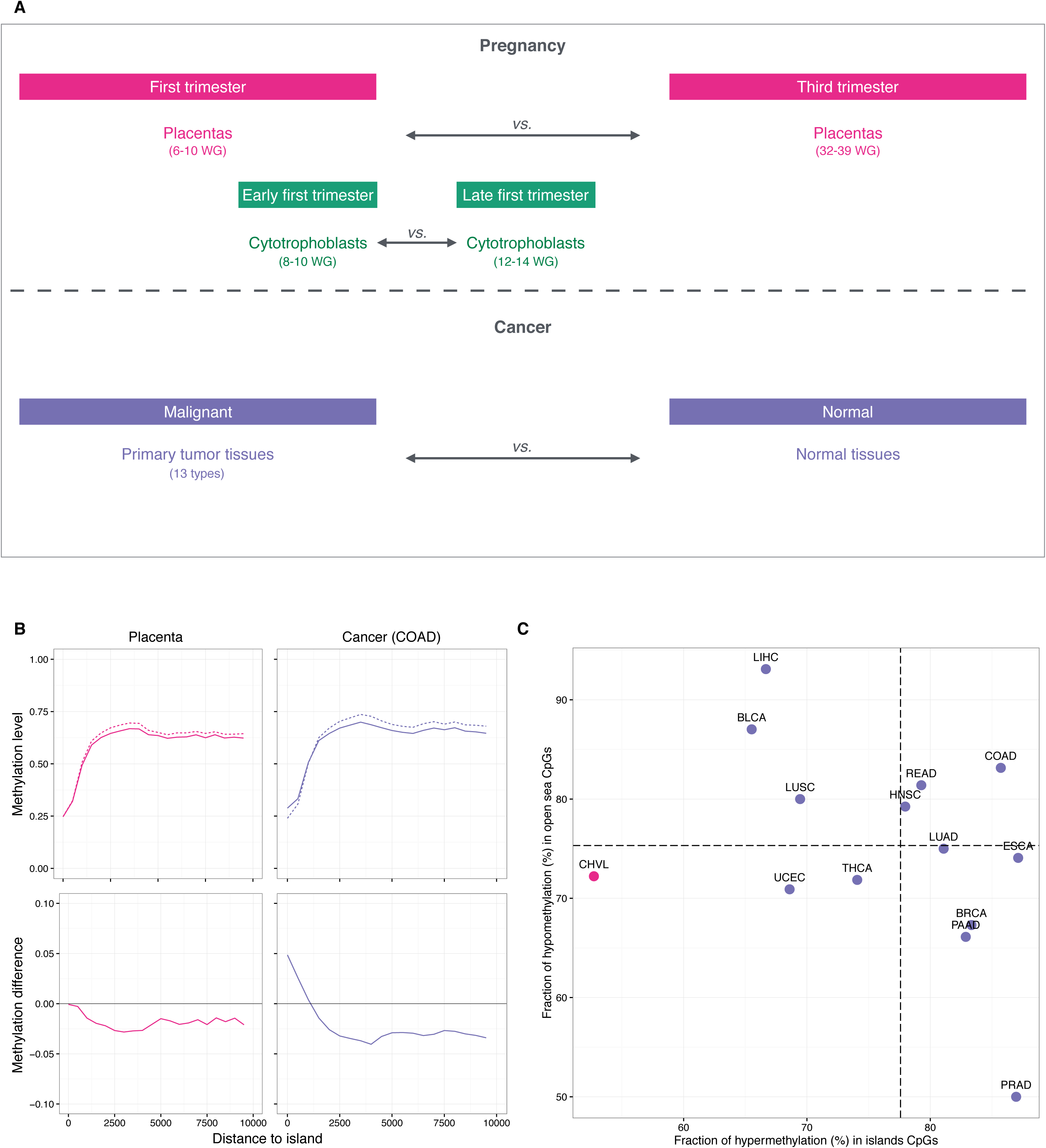
Comparison of DNA methylation in placenta and cancer. **(a)** Design of the study. **(b)** Top: methylation level plotted against the distance to the nearest CpG island. In pink, the solid line denotes first trimester samples and the dashed line denotes third trimester chorionic villi samples. In purple, the solid line denotes colon adenocarcinoma tumor samples and the dashed line denotes corresponding normal tissue samples. Bottom: methylation difference in first *vs*. third trimester chorionic villi samples (pink) and tumor *vs.* normal tissue samples (purple). **(c)** Fraction of hypermethylation (hypermethylation alterations / total alterations) in island CpGs (x-axis) *vs*. fraction of hypomethylation (hypomethylation alterations / total alterations) in open sea CpGs (y-axis) in placenta (pink) and cancer samples (purple). CpGs were classified as altered if they displayed an absolute methylation difference > 0.05 with a FDR q-value < 0.05. Horizontal and vertical dashed lines represent average fractions in cancer. Abbreviations as in Table 1.

### Large hypomethylated blocks distinguish first from third trimester placenta and tumors from matched normal tissues

Given the similarities between the patterns of hypomethylation in first trimester placental chorionic villi and cancers, we sought to further characterize the genomic regions involved. Methylomes of solid tumors have been shown to display hypomethylated blocks, defined as large regions within which the average methylation is reduced compared to normal tissues [8, 9]. To search for such hypomethylated blocks, we used the block-finding procedure implemented in the R/Bioconductor *minfi* package (See Methods, [13, 14]). We identified 1,240 blocks of differential methylation distinguishing first from third trimester placental chorionic villi samples, spanning 345 Mb (approximately 12% of the genome). Among these placenta blocks, 93% are hypomethylated in the first trimester (Table 1). On average, each individual placenta hypomethylated block has a length of 283 kb (median 225 kb) and contains 2 genes (Fig. 2a,c,d). We investigated the similarity of placenta hypomethylated blocks to those observed in cancer by applying the same block-finding procedure to the 13 cancer types from TCGA. Among these cancers, there are on average 1,713 blocks (965 – 2,195) of differential methylation distinguishing tumors from their corresponding normal tissue samples. Their coverage ranges from 212 Mb (approximately 7% of the genome) in prostate adenocarcinoma to 1,078 Mb (approximately 36% of the genome) in hepatocellular carcinoma. On average, 87% (42% – 100%) of these are hypomethylated (Table 1). In colon adenocarcinoma, taken as an illustrative example, the average hypomethylated block has a length of 493 kb (median 376 kb) and contains 4 genes (Fig. 2b,c,d). Interestingly, the hypomethylated blocks found in cancers overlapped on average 43% of placenta hypomethylated blocks (> 5 kb shared), with statistically significant co-localization (Bonferroni-adjusted alpha = 0.05) in a subset of five tumor types; namely, bladder urothelial carcinoma, colon adenocarcinoma, head and neck squamous cell carcinoma, pancreatic adenocarcinoma, and rectum adenocarcinoma (Figure S3). Taken together, these observations reveal that the patterns of hypomethylation that distinguish first from third trimester placental tissues are similar in structure, size and location to those that distinguish tumors from matched normal tissues.

**Fig 2.**
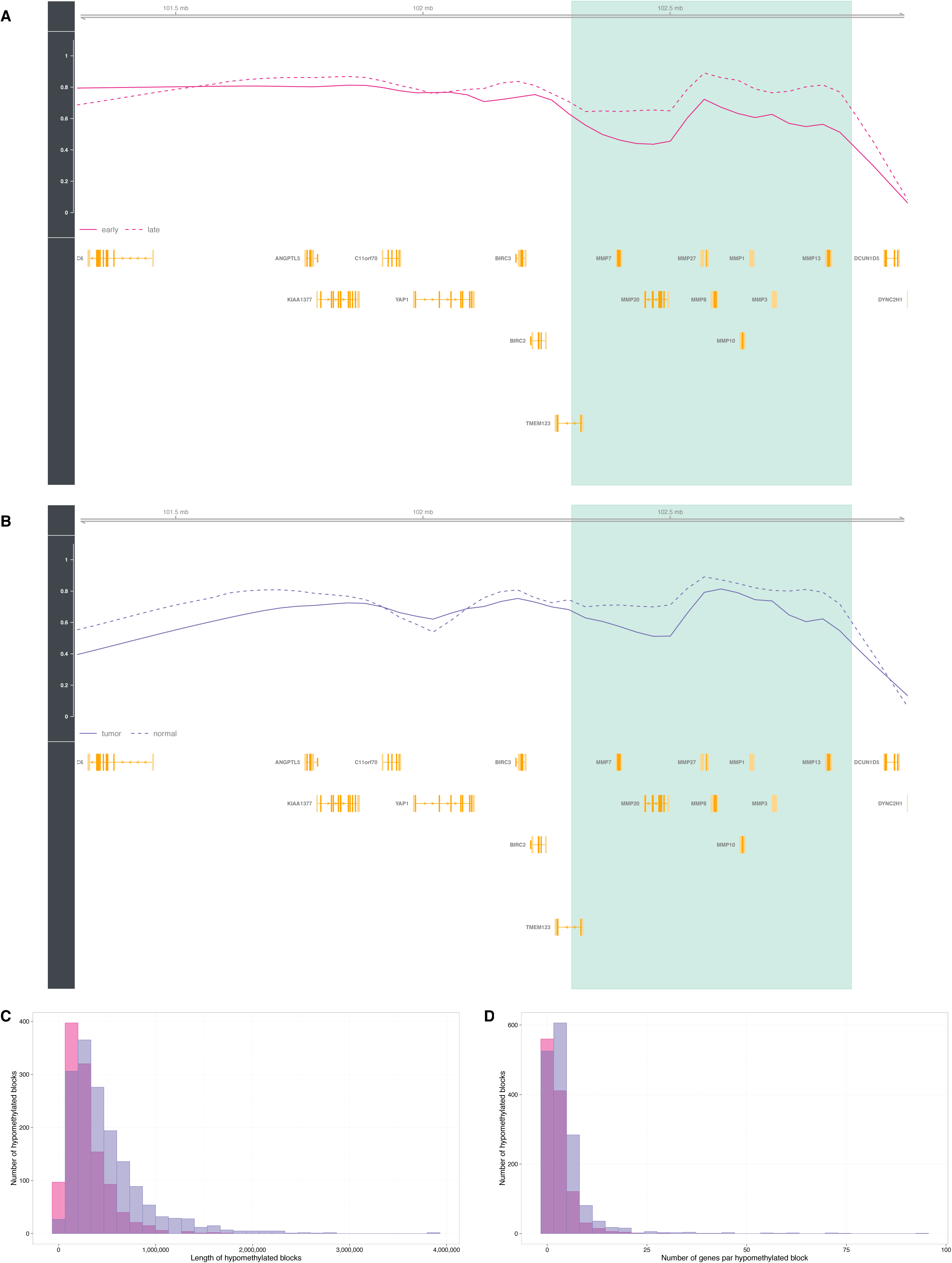
Hypomethylated blocks in placenta and cancer. **(a)** The light green zone shows an illustrative placenta hypomethylated block representing a difference in average methylation between first trimester (solid line) and third trimester (dashed line) chorionic villi samples. Gene loci are marked in yellow. **(b)** The light green zone shows an illustrative cancer hypomethylated block representing a difference in average methylation between colon adenocarcinoma tumor (solid line) and corresponding normal tissue (dashed line) samples. Gene loci are marked in yellow. **(c)** Distributions of the lengths of placenta hypomethylated blocks in pink and cancer hypomethylated blocks (colon adenocarcinoma) in purple. **(d)** Distributions of the number of genes per placenta hypomethylated blocks in pink, and cancer hypomethylated blocks in purple (colon adenocarcinoma). Abbreviations as in Table 1.

**Table 1.** Large blocks of differential methylation in placenta and cancer tissues.

|  | Blocks (N) | Coverage (Mb) | Hypo (%) |
| --- | --- | --- | --- |
| **First trimester vs. Third trimester** |  |  |  |
| Chorionic villi (CHVL) | 1240 | 345.65 | 93 |
| **Malignant vs. Normal** |  |  |  |
| Bladder urothelial carcinoma (BLCA) | 1865 | 886.54 | 99 |
| Breast invasive carcinoma (BRCA) | 2195 | 773.95 | 74 |
| Colon adenocarcinoma (COAD) | 1978 | 843.48 | 95 |
| Esophageal carcinoma (ESCA) | 965 | 305.96 | 94 |
| Head & neck squamous cell carcinoma (HNSC) | 2099 | 916.08 | 89 |
| Liver hepatocellular carcinoma (LIHC) | 1856 | 1078.46 | 100 |
| Lung adenocarcinoma (LUAD) | 1620 | 593.05 | 90 |
| Lung squamous cell carcinoma (LUSC) | 2063 | 965.50 | 93 |
| Pancreatic adenocarcinoma (PAAD) | 1058 | 211.53 | 91 |
| Prostate adenocarcinoma (PRAD) | 2105 | 567.79 | 42 |
| Rectum adenocarcinoma (READ) | 1183 | 421.84 | 100 |
| Thyroid carcinoma (THCA) | 1752 | 421.71 | 84 |
| Uterine corpus endometrioid carcinoma (UCEC) | 2041 | 686.13 | 79 |

### Hypomethylated blocks also distinguish placental villous cytotrophoblasts before and after they come into contact with maternal blood

Since substantial environment and cell type composition changes in the placenta occur during the nine months of gestation, we next sought to more narrowly define the time period and cell types associated with the observed placental methylome changes. We hypothesized that significant changes may occur concurrently with the major physiological shifts that occur when maternal blood first comes into contact with the chorionic villi during the 10th to 12th weeks of pregnancy [11]. To investigate methylation changes that occur during this period, we looked for differences that arise between the early first trimester (8-10 WG, n = 9) and late first trimester (12-14 WG, n = 10) villous cytotrophoblast samples (Fig 1a; Table S1), isolated *ex vivo* from chorionic villi samples using sequential enzymatic digestions. DNA methylation was profiled using the 450k array. At the individual probe level, we found that, as in the chorionic villi samples, CpGs distal from islands showed hypomethylation in the early first trimester villous cytotrophoblasts, while there were almost no methylation changes at CpG islands (Table S2 and Figure S4). Then, using the same block-finding procedure, we also identified large regions of differential methylation distinguishing early from late first trimester villous cytotrophoblast samples. There were 997 blocks spanning 223 Mb (approximately 7% of their genome), of which 79% are hypomethylated. On average, each individual cytotrophoblast hypomethylated block has a length of 261 kb (median 219 kb) and contains 2 genes (Fig. 3a). Importantly, cytotrophoblast hypomethylated blocks have a highly significant 85% overlap (p < 0.001) with hypomethylated blocks identified in bulk placental chorionic villi samples (Fig. 3b). This finding indicates that the placenta hypomethylated blocks identified are a phenomenon observable in individual cell types, rather than a consequence of cell type composition changes during pregnancy.

**Fig 3.**
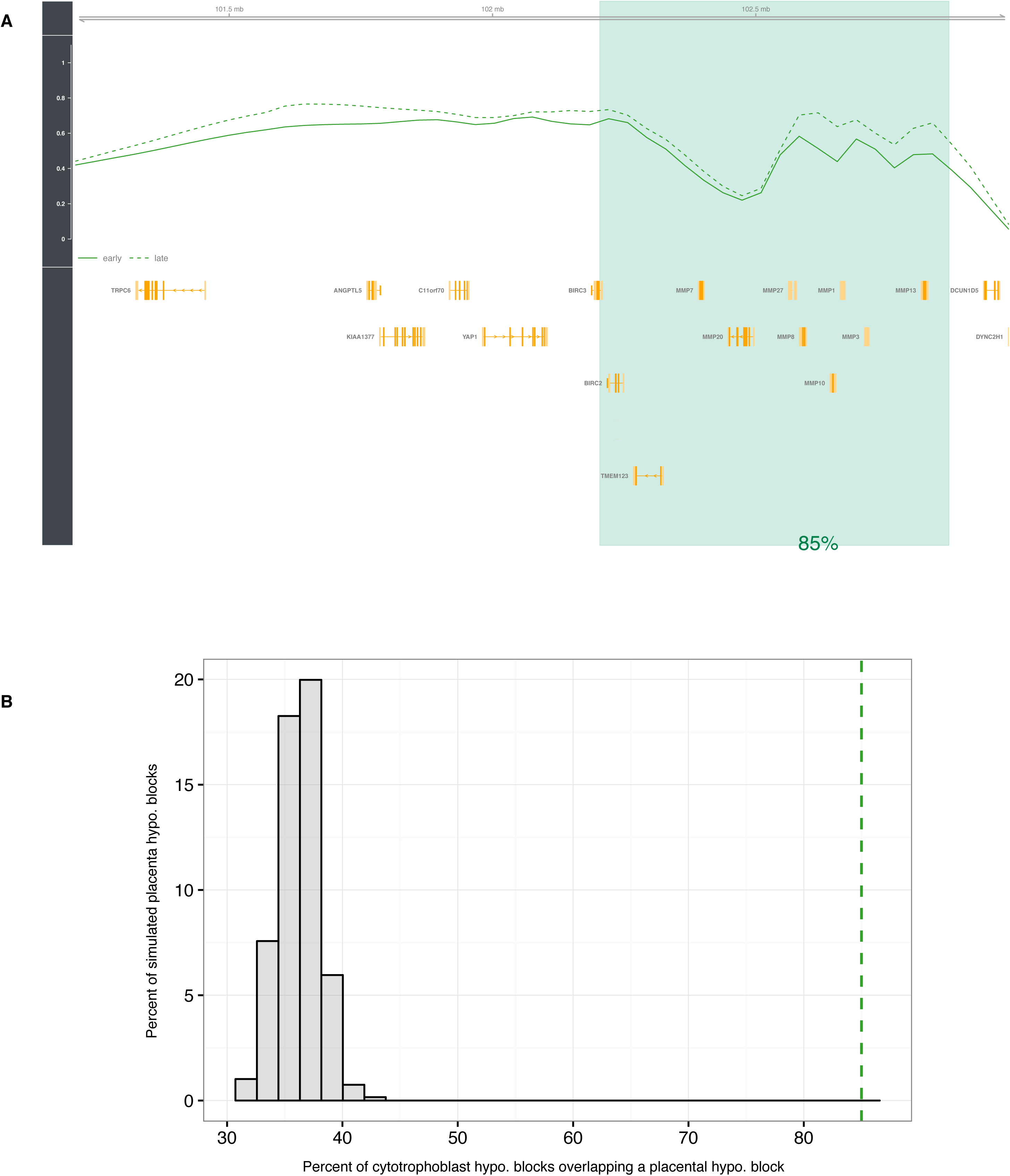
Hypomethylated blocks in isolated villous cytotrophoblasts. **(a)** The light green zone shows an illustrative cytotrophoblast hypomethylated block representing a difference in average methylation between early first trimester (solid line) and late first trimester (dashed line) villous cytotrophoblast samples. Gene loci are marked in yellow. **(b)** Null distribution of overlap fraction between cytotrophoblast hypomethylated blocks and randomly sampled regions equal in size to placenta hypomethylated blocks (chorionic villi). The dashed green line represents the observed overlap fraction (85%, p-value < 0.001). Abbreviations as in Table 1.

### Cancer and placenta hypomethylated blocks encompass genes related to hallmark cancer pathways

To gain insight into the biological functions potentially affected by hypomethylated blocks, we performed gene set analyses using MsigDB hallmark gene sets [15], and Ingenuity Pathway Analysis (IPA) gene sets related to immune response (Fig 4a; Figure S5). Gene set enrichment was assessed for genes encompassed in: (*i*) placenta and cytotrophoblast hypomethylated blocks; (*ii*) cancer hypomethylated blocks (using colon adenocarcinoma as an illustrative example of cancers); (*iii*) cancer/placenta hypomethylated blocks, i.e. cancer hypomethylated blocks that co-localize with placenta hypomethylated blocks; and (*iv*) cancer unique hypomethylated blocks, i.e. cancer hypomethylated blocks that do not co-localize with placenta hypomethylated blocks. As expected, placenta and cytotrophoblast hypomethylated blocks showed similar gene set enrichment (*p*-value < 0.05), including “*Allograft rejection*”, “*Inflammatory response*” and “*EMT*To investigate potentially shared epigenetic mechanisms between cancer and placenta, we focused on the subset of cancer/placenta hypomethylated blocks. This 44% subset of cancer hypomethylated blocks is enriched for 8 out 9 gene sets significant in the full set, including “*Allograft rejectiori*”, “*Angiogenesis*” and “*EMT*”. Interestingly, we also found two gene sets that are significant only when looking specifically at cancer/placenta hypomethylated blocks, namely “*TNF-alpha signaling via NF-kB*” (p-value = 0.005) and “*Down-regulated UV response*” (p-value = 0.01). We finally investigated the expression profile of the corresponding genes located within cancer/placenta hypomethylated blocks using RNA-sequencing data from TCGA for colon adenocarcinoma. An IPA analysis revealed two networks of genes that are differentially expressed in tumor *vs.* normal tissues (FDR q-value < 0.01). The first network was based on “*TNF-alpha signaling via NF-kB*” and included: *CD83*, *CSF2*, *CXCL2*, *CXCL3*, *DUSP4*,*INHBA*, *OLR1* and *SERPINB2*, which were significantly up-regulated; and *CD69*, *CCL5* and *KLF6* which were significantly down-regulated (Fig. 4b). The second network was based on “Down-regulated UV response” and included: *ANXA2*, *BDNF* and *COL3A1*, which were significantly up-regulated; and *AMPH*, *EFEMP1*, *ITGB3*, *KIT*, *MAGI2*, *NRP1* and *SNAI2,* which were significantly down-regulated (Fig. 4c). Notably, the up-regulated NF-kB complex represents a key hub in both networks. These results highlight processes with potentially shared epigenetic regulatory mechanisms during carcinogenesis and placentogenesis.

**Fig 4.**
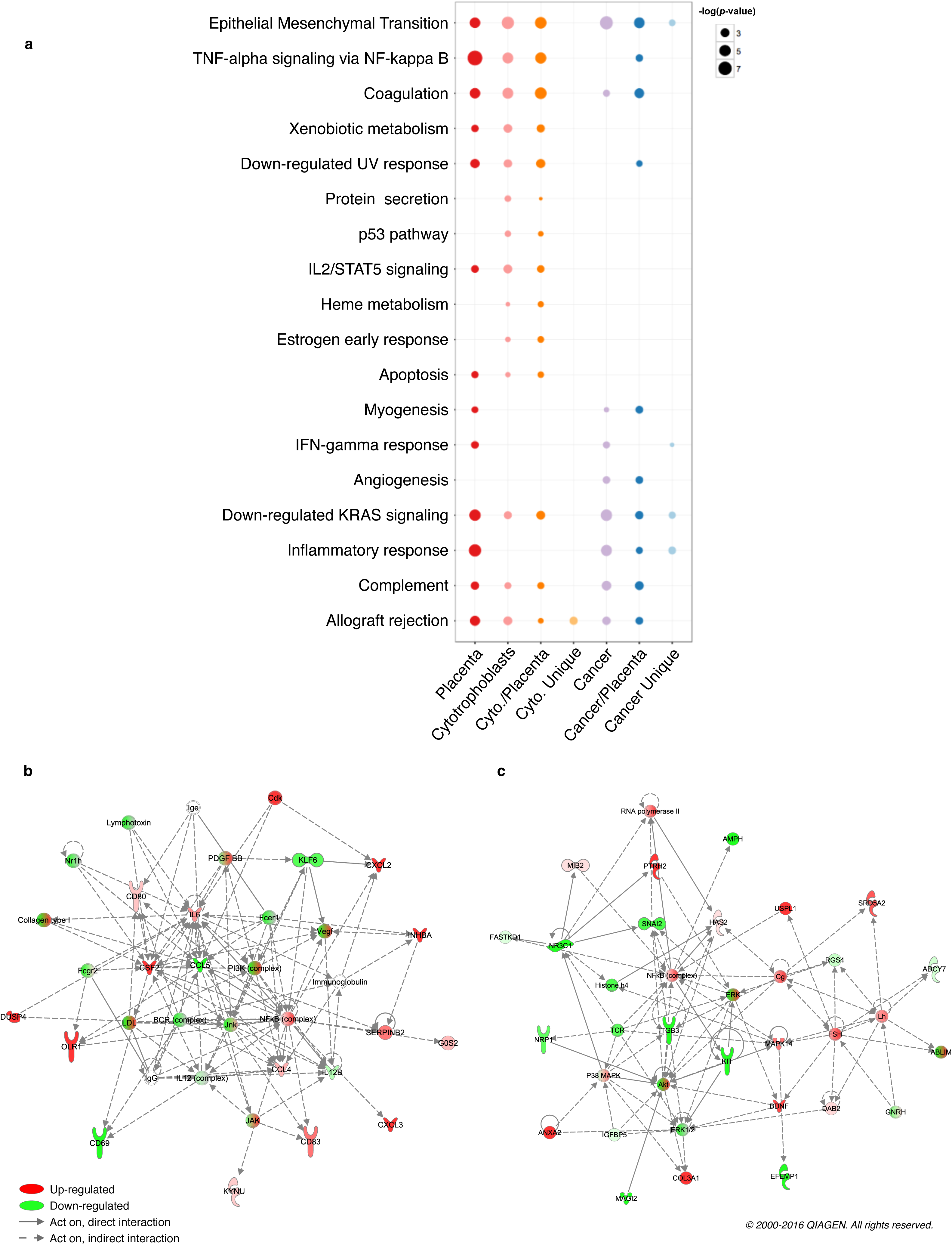
Gene set enrichment for genes encompassed in placenta and cancer hypomethylated blocks. **(a)** Bubble chart representing the enrichment of MsigDB hallmark gene sets in placenta and cancer hypomethylated blocks (colon adenocarcinoma). “Cancer/Placenta” (“Cancer Unique”) represents genes in cancer hypomethylated blocks that co-localize (do not co-localize) with a placenta hypomethylated block. Only the significant enrichments (p-value < 0.05) are presented. Networks of genes belonging to cancer/placenta blocks and relating to the **(b)** “TNF-alpha signaling via NF-kappa B” gene set and the **(c)** “UV Response - Down” gene set were identified using IPA software. Abbreviations as in Table 1.

## DISCUSSION & CONCLUSIONS

Over the 20th century, successive technological innovations have allowed the characterization of striking similarities between placental and cancer cells. The microscope initially allowed the observation of common phenotypes [16, 17], and later modern biochemistry and molecular biology allowed the characterization of molecular circuits underlying these similarities: thus, we now know that placental cells, even though they do not possesses the somatically altered genomes characteristic of cancer cells, rely on hallmark cancer molecular programs to establish pregnancy [1–4, 18]. More recently, independent studies in cancer biology and placenta biology have suggested similarities between epigenetic patterns in placental and cancer cells [5, 6].

The study presented here focused on the identification of commonalities between methylation dynamics during pregnancy and neoplastic transformation. We conducted the first direct genome-wide comparison between DNA methylation changes in placenta throughout pregnancy, and in cancer during neoplastic transformation. We used publically available DNA methylation and gene expression data, and complemented this with a newly generated dataset representing the first genome-wide DNA methylation profiles from isolated villous cytotrophoblasts obtained before and after the cells come into contact with maternal blood, a key step in the establishment of pregnancy. We report here that epigenetic similarities between cancer and placenta are most striking in early pregnancy placenta, and that the similarities are partially erased as pregnancy progresses. We also find that these shared epigenetic patterns relate to pivotal genes in both placentogenesis and carcinogenesis.

We first found that placental methylome remodeling during pregnancy involves methylation gains distal from CpG islands. The affected regions largely overlap with those that become hypomethylated during neoplastic transformation. We also observed that CpG islands remain largely unaltered during pregnancy, unlike in cancer where these regions gain methylation. Our findings are consistent with results from a high performance liquid chromatography (HPLC) assay that shows an increase in methylation levels during pregnancy [19]. These results suggest that the hypermethylated state of tumor suppressor promoters, including *RASSF1A*, *APC*, *SFRP2*, *WIF1*, and *EN1*, described by previous studies on DNA methylation in the placenta [5, 20–22] remains largely stable throughout pregnancy. It also suggests two mechanisms underlying DNA methylation dynamics: one mechanism common to the placenta during pregnancy and tumors during neoplastic transformation; and another mechanism of DNA methylation specific to tumors.

We found that hypomethylation in first relative to third trimester chorionic villi is organized into large blocks with an average length of 283 kb. In 2013, Schroeder *et al*. first described large-scale hypomethylated regions in chorionic villi [23], which thus represents the first known normal tissue showing clear evidence of large-scale contiguous blocks of hypomethylation of the kind previously been described in cancer. This observation was in line with the historic observation of a global hypomethylation characteristic of placenta and cancers compared to other tissues [24, 25]. Hypomethylated blocks have initially been described in cancers using whole-genome bisulfite sequencing: first, in colon tumors [8, 26]; then, in breast cancer cell line [27], medulloblastoma tumors [28] and in EBV immortalized B-cells [29]. In 2014, Timp *et al*. reported the first integrated genome-wide analysis of DNA methylation of six different tumor types (breast, colon, lung, pancreas adenocarcinoma, pancreas neuroendocrine tumors, thyroid cancer samples) using the 450k array [9]. These studies have thus established hypomethylated blocks as an universal defining epigenetic alteration in human solid tumors. Using TCGA data for 13 cancer types, we also observed hypomethylated blocks in breast, colon, lung, pancreas and thyroid cancers, as well as described such hypomethylation patterns for the first time in liver, head and neck, bladder, uterus, esophageal and prostate cancers. Strikingly, in most cases, the number of blocks, size and overall hypomethylation we found in cancers were similar to the ones we found in chorionic villi. Cancer and chorionic villi hypomethylated blocks also display frequent co-localization, which was significant in several cancer types. Taken together, our observations provide the first evidence of dynamic change of these large-scale hypomethylated regions throughout pregnancy, and highlight a striking parallel with hypomethylation patterns in cancers.

Moreover, we also observed that hypomethylated blocks distinguish villous cytotrophoblasts before and after they first come into contact with maternal blood (weeks 10-12), providing the first evidence of an isolated non-cancer cell type with large-scale hypomethylated blocks. Furthermore, the result indicates that our finding of hypomethylated blocks in the placenta is not an artifact of cell-type confounding related to changes in cellular composition during pregnancy. It also suggests that many of the methylation differences between early and late pregnancy placenta are associated with changes in the cytotrophoblast microenvironment including exposure to components of maternal blood and higher oxygen tension. This time point has been also shown to be associated with changes in proliferative, invasive and fusion abilities of the cytotrophoblasts [11, 30].

Finally, our data suggest that cancer and placenta hypomethylated blocks might be involved in the regulation of hallmark cancer pathways. The regulatory implications of megabase-scale hypomethylation is poorly understood, but the finding that it is associated with increased gene expression variability has led to the suggestion that it may provide a mechanism for a phenotypic plasticity that allows adaptation and rapid growth in the setting of a host tissue [31]. Supporting this hypothesis, we found that cancer and placenta hypomethylated blocks are enriched for pathways including EMT, allograft rejection and inflammatory response. Interestingly, we also identified two enriched gene sets that are detectable only when focusing on the subset of cancer/placenta hypomethylated blocks rather than the full set of cancer hypomethylated blocks. Differentially expressed genes in these sets are involved in predicted networks centered on NF-kB, a transcription factor complex playing a critical role in inflammation [32]. This observation is in line with the hypothesis that the establishment of pregnancy and cancer are both pro-inflammatory states [33, 34]. These networks include *BDNF*, *SERPINB2* and *CXCL3*, genes previously described as instrumental in both placentogenesis and cancerogenesis. The brain-derived neurotrophic factor (BDNF) has been linked to placental development (Kawamura et al., 2009) and it has also been found to regulate cell motility in colon cancer [35]. Plasminogen Activator Inhibitor Type 2 (PAI-2), encoded by *SERPINB2*, was initially identified in the placenta [36] and low levels of PAI-2 has been linked to preeclampsia [37]. Since then, numerous studies have attempted to elucidate its role in invasion and metastasis [38]. *CXCL3* has been implicated in placental invasion and is aberrantly expressed both in severe preeclampsia [39] and in cancer during esophageal carcinogenesis [40]. Our analyses also pointed to immune-related genes which were down-modulated in colon tumors, including *CD69* and *CCL5*. This last observation could be explained by the presence within the immune infiltrate of suppressive subpopulations such as regulatory T cells (Tregs), M2 macrophages or myeloid-derived suppressive cells (MDSC), that have previously been linked to tolerance induction both in cancer and pregnancy [2, 4, 41, 42]. Our approach thus suggests a means for identifying genes potentially regulated by similar epigenetic mechanisms during pregnancy and neoplastic transformation.

This pilot study suggests the existence of megabase-scale patterns of hypomethylation that characterize the early pregnancy placenta and tumors, and which might contribute to the regulation of molecular programs allowing cellular growth advantage at the expense of the host. Further analyses of these patterns could eventually lead to the identification of critical epigenetic switches that prevent healthy placentas from degenerating into tumors, and whose failure allows tumor development. The study demonstrates the relevance of placental research as a platform for innovative approaches in oncology as well as in obstetrics and gynecology.

## METHODS

### DNA methylation data

Villous cytotrophoblast data was generated on the 450k platform and normalized using the *preprocessFunnorm* function implemented in the *minfi* package of the R/Bioconductor software environment [13, 14]. Public 450k data for chorionic villi and cancers was obtained from GEO (<u>GSE49343</u>)[10] and TCGA (http://cancergenome.nih.gov/), respectively (**Supplementary Table 1**). The calculated Beta values from GEO were downloaded using the *getGenomicRatioSetFromGEO* function of *minfi* [13, 14], and the calculated Beta values from TCGA (level 2 data, Feb 24th 2015 version) were downloaded via the UCSC Cancer Genomics Browser website (https://genome-cancer.ucsc.edu/). The solid tumor *vs.* normal tissue samples spanned 13 cancer types: bladder urothelial carcinoma (BLCA); breast invasive carcinoma (BRCA); colon adenocarcinoma (COAD); esophageal carcinoma (ESCA); head and neck squamous cell carcinoma (HNSC); liver hepatocellular carcinoma (LIHC); lung adenocarcinoma (LUAD); lung squamous cell carcinoma (LUSC); pancreatic adenocarcinoma (PAAD); prostate adenocarcinoma (PRAD); rectum adenocarcinoma (READ); thyroid carcinoma (THCA); and uterine corpus endometrioid carcinoma (UCEC) (Supplementary Table 1).

### Isolation and purification of human villous cytotrophoblasts

The *ex vivo* villous cytotrophoblast samples were isolated from placental tissues (chorionic villi) and extemporaneously frozen. Sequential enzymatic digestion (Trypsine and DNAse) based on methods previously described by Kliman *et al*. [43] were used with modifications to isolate villous cytotrophoblast cells [11]. The isolation step was followed by a purification step based on Percoll gradient fractionation. A subset of the purified villous cytotrophoblasts were characterized by immunolabeling using the cytokeratin 7 (CK7) and their ability to aggregate after 48 hours and to form syncytiotrophoblasts at 72 hours of culture. DNA was extracted using the GenElute™ Mammalian Genomic DNA Miniprep Kit (Sigma-Aldrich).

### Probe level analysis

For each probe, we averaged the methylation level (Beta value between 0 and 1) across individuals in each case group (e.g. chorionic villi first trimester or colon adenocarcinoma tumor) and control group (e.g. chorionic villi third trimester or colon adenocarcinoma normal). We then computed the difference at the probe level between these case and control groups in placenta and cancer. Student’s t-test was used to assess statistical significance of differences, and the false discovery rate procedure implemented in the R/Bioconductor *qvalue* package was used to correct for multiple testing [44, 45]. Methylation differences were termed biologically meaningful when above 0.05 and statistically significant for an FDR q-value below 0.05.

### Block-finding procedure

In order to investigate blocks (megabase-scale changes in methylation) in Illumina 450k data, we used the block-finding procedure implemented in the R/Bioconductor *minfi* package [13, 14]. This procedure uses open-sea CpGs, i.e. those distal from CpG islands, to identify large-scale methylation patterns. The 485,512 probes on this array target regions including CpG islands (31% of the array), CpG island shores (23%) and shelves (10%) located adjacent to CpG islands, while the remaining probes represent open-sea CpGs located at least 4 kb away from CpG islands (36%) [12]. Briefly, the procedure groups adjacent open sea CpGs into clusters with a maximum gap width of 500 bp. Clusters with a width > 1,500 bp are subdivided. Methylation values at CpGs within each cluster are averaged, resulting in a single mean estimate per cluster. Student’s t-statistics comparing cases (e.g. chorionic villi first trimester or colon adenocarcinoma tumor) and controls (e.g. chorionic villi third trimester or colon adenocarcinoma normal) were calculated for each cluster. Then, clusters within 250 kb from each other were grouped together. Finally, the Bumphunter algorithm [46] was applied within each group to detect blocks exhibiting differences in average methylation between cases and controls. The main steps of this algorithm are the following: (*i*) fitting a loess curve with a 250 kb smoothing window through the t-statistics; (*ii*) identifying regions with an absolute smoothed t-statistic in the top 2.5% as putative blocks, and (*iii*) determining block p-values based on the likelihood of such dimensions occurring by chance through a sample permutation test (1,000 permutations) using *minfi* default parameters. The p-value is calculated as the fraction of permutations that generate a block of equal or larger dimensions. Blocks containing at least five probe clusters with a q-value below 0.05 were declared biologically relevant and statistically significant. Also, as we used microarray data, and not whole-genome bisulfite sequencing data, we could only identify blocks located in regions included in the design of the 450k assay; we could not tell if regions excluded from the array contain blocks.

### Block overlap P values

For each tissue, tables gathering all characteristics of the methylation blocks we detected were converted into GRanges objects of the R/Bioconductor software environment to allow convenient manipulation of information corresponding to genomic intervals [47]. Overlaps between each set of hypomethylated blocks found in cancers (the queries) and the set of placenta hypomethylated blocks (the subject) were assessed as the fraction of cancer hypomethylated blocks that overlapped a placenta hypomethylated block by at least 5 kb. Significance of the overlap rate was determined using Monte Carlo simulations. We generated 1, 000 random lists of simulated placenta hypomethylated blocks. We constrained the simulation such that, compared to observed placenta hypomethylated blocks, the simulated placenta hypomethylated blocks had: the same genomic coverage (within 10%) and the same number of collapsed open sea probes. We further required that hypomethylated blocks within a simulated set were non-overlapping by resampling until non-overlapping lists were obtained. Then, we counted overlaps between each of list of simulated subject (placenta) hypomethylated blocks and the observed query lists (each cancer). The 1,000 overlap fractions formed the null distribution of overlaps obtained by chance. We then compared these null distributions with the observed overlap fractions. To account for multiple testing, we declared significant those overlaps with a p-value less than the Bonferroni-adjusted threshold i.e. p-value/number of comparisons = 0.05/13. Notably, this approach avoids biases due to the design of the 450k microarray platform.

### MsigDB and IPA gene set enrichment analyses

The gene universe was defined as unique gene symbols in the R/Bioconductor *Homo sapiens* package with a TSS within a region probed by block-finding procedure (i.e. regions defined by the *cpgCollapse* function of *minfi*). Using the R/Bioconductor *GeneOverlap* package, gene set enrichment p-values for MsigDB hallmark gene sets [15] and IPA (http://www.ingenuity.com/) gene sets related to immune response were computed for genes encompassed in: (*i*) placenta and cytotrophoblast hypomethylated blocks; (*ii*) cancer hypomethylated blocks (using colon adenocarcinoma as an illustrative example of cancers); (*iii*) cancer/placenta hypomethylated blocks, i.e. cancer hypomethylated blocks that co-localize with placenta hypomethylated blocks; and (*iv*) cancer unique hypomethylated blocks, i.e. cancer hypomethylated blocks that do not co-localize with placenta hypomethylated blocks. Rows of Fig. 4a were ordered using the *ward.2* method for hierarchical clustering built in the R software environment. Gene expression profiles of colon adenocarcinoma primary tumors (n = 241) and normal tissues (n = 39) measured by TCGA using Illumina HiSeq 2000 RNA-Seq platform were downloaded via the UCSC Cancer Genomics Browser website (level 3 RNA-seq profiles, RNAseqV2 normalized RSEM, Feb 24th 2015 version). The differentially expressed gene list was produced using the R/Bioconductor *limma* package [48].

## LIST OF ABBREVIATIONS

450k: Illumina HumanMethylation450
BLCA: bladder urothelial carcinoma
BRCA: breast invasive carcinoma
COAD: colon adenocarcinoma
EMT: epithelial-mesenchymal transition
ESCA: Esophageal carcinoma
FDR: false discovery rate
GEO: Gene Expression Omnibus
HNSC: head and neck squamous cell carcinoma
IPA: Ingenuity Pathway Analysis
LIHC: liver hepatocellular carcinoma
LUAD: lung adenocarcinoma
LUSC: lung squamous cell carcinoma
NF-kB: nuclear factor kappa B
PAAD: pancreatic adenocarcinoma
PRAD: prostate adenocarcinoma
READ: rectum adenocarcinoma
THCA: thyroid carcinoma
TCGA: The Cancer Genome Atlas
UCEC: uterine corpus endometrial carcinoma
WG: weeks of gestation

## DECLARATIONS

### Ethics approval and consent to participate

The *ex vivo* villous cytotrophoblast samples were obtained following legal voluntary interruption of pregnancy in patients at the Department of Obstetrics and Gynecology at Cochin Port-Royal Hospital (Paris, France). Patients gave their informed written consent and the local ethics committee (CCP IDF1, N°13909, Paris, France) approved the study.

### Consent for publication

Not applicable.

### Availability of data and materials

DNA methylation profiles (Illumina 450k data) of the first trimester villous cytotrophoblasts are available in GEO under accession number GSE93208.

### Competing interests

The authors declare that they have no competing interests.

### Funding

This work was supported by: the Association pour la recherche en cancerologie de Saint-Cloud (ARCS); ADEBIOPHARM; Genevieve and Jean-Paul Driot Transformative Research Grant; Ouidad Hachem Foundation Grant; Philippe, Laurent and Stéphanie Bloch Cancer Research Grant; Sally Paget-Brown Translational Research Grant; and French state funds within the Investissements d’Avenir program (ANR-11-IDEX-0004-02).

### Authors’ contributions

A.V.N. and M.J.A. conceived and directed the study. A.V.N., D.N.B. and M.J.A. analyzed and interpreted the data with the support of D.K., D.B., V.D.M. and T.F. S.R. preprocessed the villous cytotrophoblast samples. D.K and D.B. provided software. T.F. provided the villous cytotrophoblast samples. A.V.N. and M.J.A. wrote the manuscript. All authors read and approved the final manuscript.

## Acknowledgements

We are grateful to Raphaël Bilgraer for his support and feedback. We also thank Sophie Gil and Françoise Vibert for their help in isolating villous cytotrophoblasts.

